# Viral competence data improves rodent reservoir predictions for American orthohantaviruses

**DOI:** 10.1101/2021.01.01.425052

**Authors:** Nathaniel Mull, Colin J. Carlson, Kristian M. Forbes, Daniel J. Becker

## Abstract

Identifying reservoir host species is crucial for understanding the risk of pathogen spillover from wildlife to people. Orthohantaviruses are zoonotic pathogens primarily carried by rodents that cause the diseases hemorrhagic fever with renal syndrome (HFRS) and hantavirus cardiopulmonary syndrome (HCPS) in humans. Given their diversity and abundance, many orthohantaviruses are expected to be undiscovered, and several host relationships remain unclear, particularly in the Americas. Despite the increasing use of predictive models for understanding zoonotic reservoirs, explicit comparisons between different evidence types for demonstrating host associations, and relevance to model performance in applied settings, have not been previously made. Using multiple machine learning methods, we identified phylogenetic patterns in and predicted unidentified reservoir hosts of New World orthohantaviruses based on evidence of infection (RT-PCR data) and competence (live virus isolation data). Infection data were driven by phylogeny, unlike competence data, and boosted regression tree (BRT) models using competence data displayed higher accuracy and a narrower list of predicted reservoirs than those using infection data. Eight species were identified by both BRT models as likely orthohantavirus hosts, with a total of 98 species identified by our infection models and 14 species identified by our competence models. Hosts predicted by competence models are concentrated in the northeastern United States (particularly *Myodes gapperi* and *Reithrodontomys megalotis*) and northern South America (several members of tribe Oryzomyini) and should be key targets for empirical monitoring. More broadly, these results demonstrate the value of infection competence data for predictive models of zoonotic pathogen hosts, which can be applied across a range of settings and host-pathogen systems.

**Author Summary:** Human diseases with wildlife origins constitute a significant risk for human health. Orthohantaviruses are viruses found primarily in rodents that cause disease with high rates of mortality and other complications in humans. An important step in disease prevention is to identify which rodent species carry and transmit orthohantaviruses. By incorporating species relatedness and evidence of different levels of host capacity to be infected and transmit virus, we used predictive modeling to determine unidentified rodent hosts of orthohantaviruses. Models using host competence data outperformed models using host infection data, highlighting the importance of stronger data in model optimization. Our results highlighted roughly a dozen key target species to be monitored that are concentrated in two geographic regions—northeastern United States and northern South America. More broadly, the approaches used in this study can be applied to a variety of other host-pathogen systems that threaten public health.

## Introduction

Identifying reservoir host species (those that maintain and transmit a particular pathogen; Haydon et al. 2002) is crucial for understanding the risk of pathogen spillover from wildlife to people (zoonotic transmission; Viana et al. 2014; Plowright et al. 2017). By elucidating possible sources of zoonotic exposure, targeted strategies can be implemented to prevent or at least mitigate spillover risk. Large-scale surveillance of wildlife, often involving non-targeted sampling of a large diversity and abundance of animals, is commonly conducted shortly after disease outbreaks to search for the pathogen reservoirs (e.g., Leroy et al. 2005; Poon et al. 2005). Such studies are often expensive, time-consuming, and inefficient, particularly when there is little information to direct sampling effort (e.g., Johara et al. 2001; Poon et al. 2005; Pourrut et al. 2009). Therefore, it is imperative to develop efficient methods for identifying reservoir hosts.

Recent advances in trait-based models have increased precision, and in turn predictive power, to facilitate identification of unknown reservoirs of viruses in nature (Becker et al. 2020a; Crowley et al. 2020). However, significant questions remain about how modelers should implement these approaches, particularly in regards to the type and level of evidence for virus infection and ability for onward transmission (Becker et al. 2020b; Worsley-Tonks et al. 2020). Most models are based on serology data (i.e., antibodies), which tend to be abundant due to its relative ease and cost-effectiveness to collect. However, such information often only provides evidence of virus exposure, not necessarily current infection (Gilbert et al. 2013). Polymerase chain reaction (PCR), on the other hand, provides stronger evidence of current infection, and is a better predictor of host competence than serology data (Tolsá et al. 2018). However, virus infection does not necessarily equate to onward transmission potential. Instead, the “gold standard” and least common evidence for competent reservoir hosts (i.e., those capable of transmitting virus) is live virus isolation (Corona et al. 2018). Current understanding of how these different types of evidence alter predictive capacity is limited, despite clear differences in host associations and relevance to model performance in applied settings (i.e., future efforts to search for reservoirs).

Orthohantaviruses (Bunyavirales: Hantaviridae) are an ideal virus group to examine using predictive models, due to their broad implications for human health as zoonotic pathogens, the predicted large number of unidentified viruses (Vaheri et al. 2008), and the varying types of virus infection and competence evidence currently available from wildlife surveys. There are currently 58 described orthohantaviruses, primarily found in rodents (Laenen et al. 2019), that cause two main human diseases: hemorrhagic fever with renal syndrome (HFRS, which is common throughout the Old World) and hantavirus cardiopulmonary syndrome (HCPS or HPS, which is common throughout the New World). Because each human case is thought to be an independent spillover event from an infected rodent (Forbes et al. 2018; Avšič-Županc et al. 2019), identifying orthohantavirus reservoir host species is critical for efforts to mitigate human disease.

With few exceptions, including mole- and shrew-borne orthohantaviruses (Arai et al. 2007; Arai et al. 2008; Kang et al. 2009; Kang et al. 2011), most known orthohantaviruses infect rodents in the families Cricetidae and Muridae (superfamily Muroidea), including all orthohantaviruses that cause disease in humans (Forbes et al. 2018). Because spillover is constrained by phylogenetic distance (Streicker et al. 2010), undiscovered orthohantaviruses are also likely to be found among muroid rodents. Additionally, although the majority of described American orthohantaviruses cause disease in humans (13/22), knowledge of host relationships is weak for these viruses, and frequent discovery of novel orthohantaviruses indicates a high likelihood of unknown viruses in this part of the world (Mull et al. 2020). Efforts to predict novel orthohantavirus reservoirs can therefore be focused within New World muroids for maximum precision and impact.

In this study, we used machine learning approaches to predict reservoir hosts of unknown American orthohantaviruses. Predictions were generated by combining muroid phylogenetic and trait data with two levels of evidence for the propensity of a species to host orthohantaviruses: (1) RT-PCR (termed infection) and (2) live virus isolation (termed competence). Model performance was compared using these two evidence types to determine the power of our various methods for identifying undiscovered orthohantavirus hosts. Finally, host predictions were incorporated with geospatial data on human exposure potential to predict geographic areas with greatest zoonotic spillover risk. Generated results will guide ongoing and future efforts to discover novel orthohantaviruses and determine virus-host relationships. More broadly, determining effective modeling approaches, specifically the role of infection versus competence data, is critical to optimizing tools for identifying and understanding potential zoonotic threats to human health and security.

## Results

### Phylogenetic patterns

Across our 601 New World muroid rodent species, 9.32% displayed evidence of orthohantavirus infection, whereas only 2% were found positive for virus isolation (Fig 1). We identified intermediate phylogenetic signal in infection (*D* = 0.81) but little phylogenetic signal in competence (*D* = 0.90). For the former, phylogenetic patterns in infection departed from both randomness (*p* < 0.001) and Brownian motion (*p* < 0.001), whereas competence departed from Brownian motion (*p* < 0.001) but not phylogenetic randomness (*p* = 0.16). Results from phylogenetic factorization were qualitatively similar. We identified two rodent clades with significantly greater propensities to have orthohantavirus infection. A subclade of the genus *Peromyscus* (*n* = 24) and the whole genus *Oligoryzomys* (*n* = 20) had 37.5% and 40% of species predicted to be capable of becoming infected, respectively, compared to 8% of the paraphyletic remainder. In contrast, our analyses identified no taxonomic patterns in competence.

**Fig 1.**
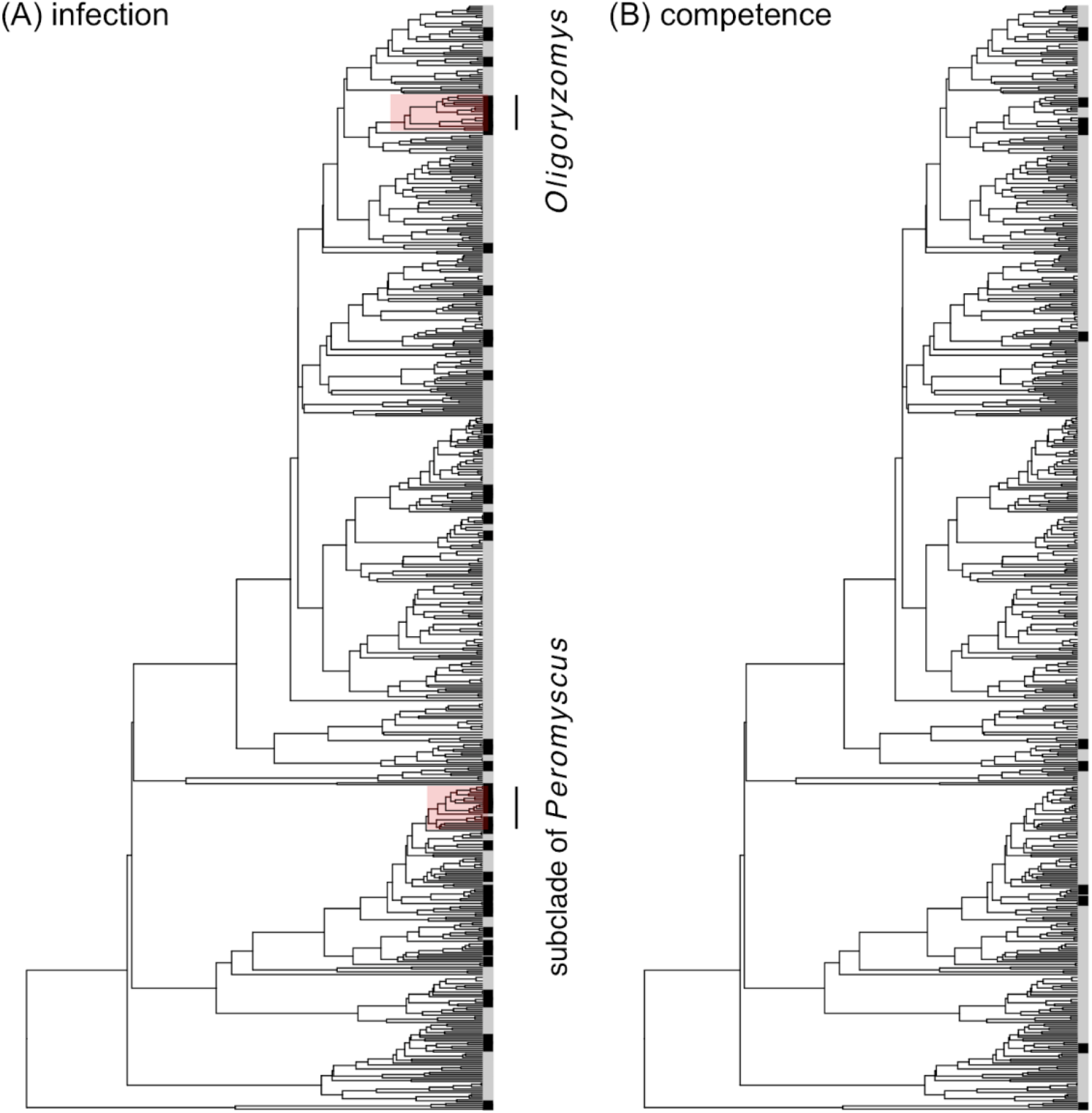
Phylogenetic distribution of orthohantavirus-positive muroid rodents in the New World. Species with evidence of infection (A, RT-PCR) or competence (B, live virus isolation) are displayed in black. Visualized in red are any clades identified through phylogenetic factorization for having greater virus positivity when compared to the paraphyletic remainder.

### Model performance

Both infection and competence BRT models distinguished orthohantavirus positive and negative rodent species with high accuracy (A*UC*= 0.92 ± 0.002). However, BRTs trained on host competence performed significantly better (A*UC*= 0.94 ± 0.003) than those trained on infection (A*UC*= 0.91 ± 0.003; *t*=6.24, *p* < 0.001; Fig 2A), resulting in a moderate effect size (*d*=0.62; Cohen 1988). Despite this difference in model performance, both models identified similar species traits as predictive of positivity (infection and competence). Rankings of variable importance were strongly correlated (ρ = 0.88, *p* < 0.001), even after removing traits with zero relative importance (*n* = 42 remaining features; ρ = 0.82, *p* < 0.001). Consistently important features for both response variables included PubMed citations, litter size, and both mammal richness and mean precipitation within the species range. Consistently unimportant features included the genera *Neotoma, Rhipidomys, Nectomys*, and *Handleyomys*. Major discrepancies included the genus *Peromyscus* and activity cycle being important predictors of infection but not competence and the genus *Oryzomys* being an important predictor of competence but not infection (Fig 2B, S2 Table). Partial dependence plots suggested that the directions of effects were largely consistent across models, with positive species being well-studied, located in mammal-rich regions, and characterized by a faster life history (S1 Fig). However, our secondary BRTs showed that citations were not predictable by traits (A*UC*= 0.49 ± 0.001), suggesting that the trait profile of positive rodents is not confounded by the traits of well-studied species.

**Fig 2.**
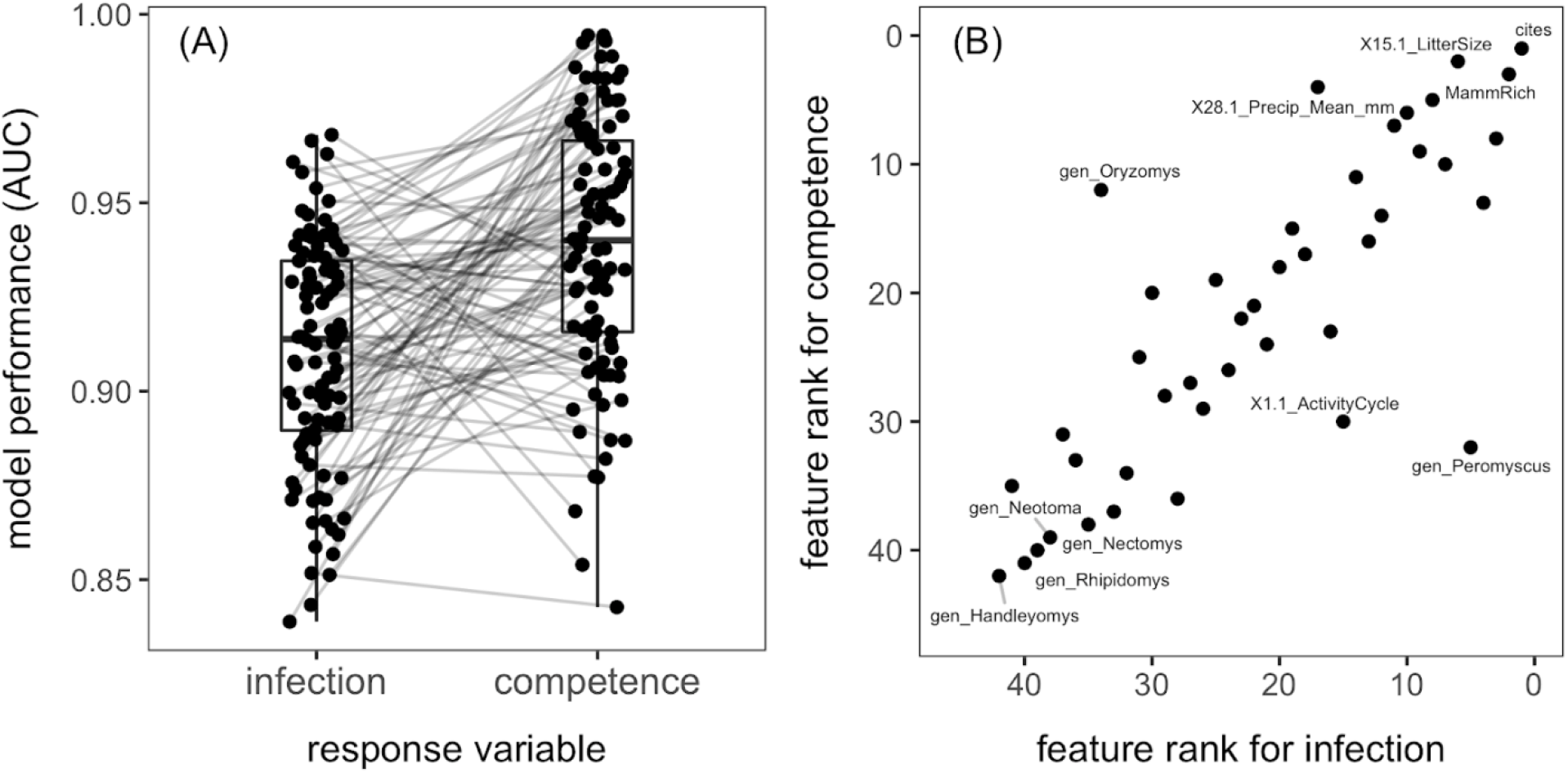
Performance of rodent orthohantavirus BRT models trained on infection versus competence data as the response. (A) Area under the receiver operating characteristic curve (AUC) across 100 random splits of training (70%) and test (30%) data. Boxplots show the median and interquartile range alongside paired AUC values. (B) Correlation between ranks of mean feature importance between models. Mean relative importance is given in S2 Table.

### Model prediction

Predicted probabilities of being an orthohantavirus host varied widely across the 601 rodent species but were only weakly positively correlated between infection and competence BRTs (ρ = 0.14, *p* < 0.001; Fig 3A). Many species with intermediate-to-high propensity scores from models based on infection had a low corresponding probability of being competent. Whereas both predictions displayed moderate phylogenetic signal (λ = 0.58 and 0.54, respectively), the taxonomic patterns identified by phylogenetic factorization largely differed between models (Fig 3B, S3 Table). For both infection and competence models, the genus *Oligoryzomys* (*n* = 20) had a greater mean probability of orthohantavirus hosting compared to the paraphyletic remainder. Predictions from infection models otherwise largely mirrored observed patterns in the data, with a subclade of the genus *Peromyscus* (*n* = 25) also having greater propensity scores 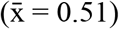, although a subclade of the genus *Oxymycterus* (*n* = 6) and the subfamily Arvicolinae (including voles, lemmings, and muskrats; *n* = 43) had greater and lower probabilities of infection (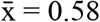 and 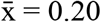, respectively). Predictions from competence models instead were clustered in the genus *Oryzomys* (*n* = 6), a subclade of *Oecomys* (*n* = 7), a smaller subclade of *Peromyscus* (*n* = 7), and the genus *Sigmodon* (*n* = 13), all of which had greater probabilities of being reservoirs.

**Fig 3.**
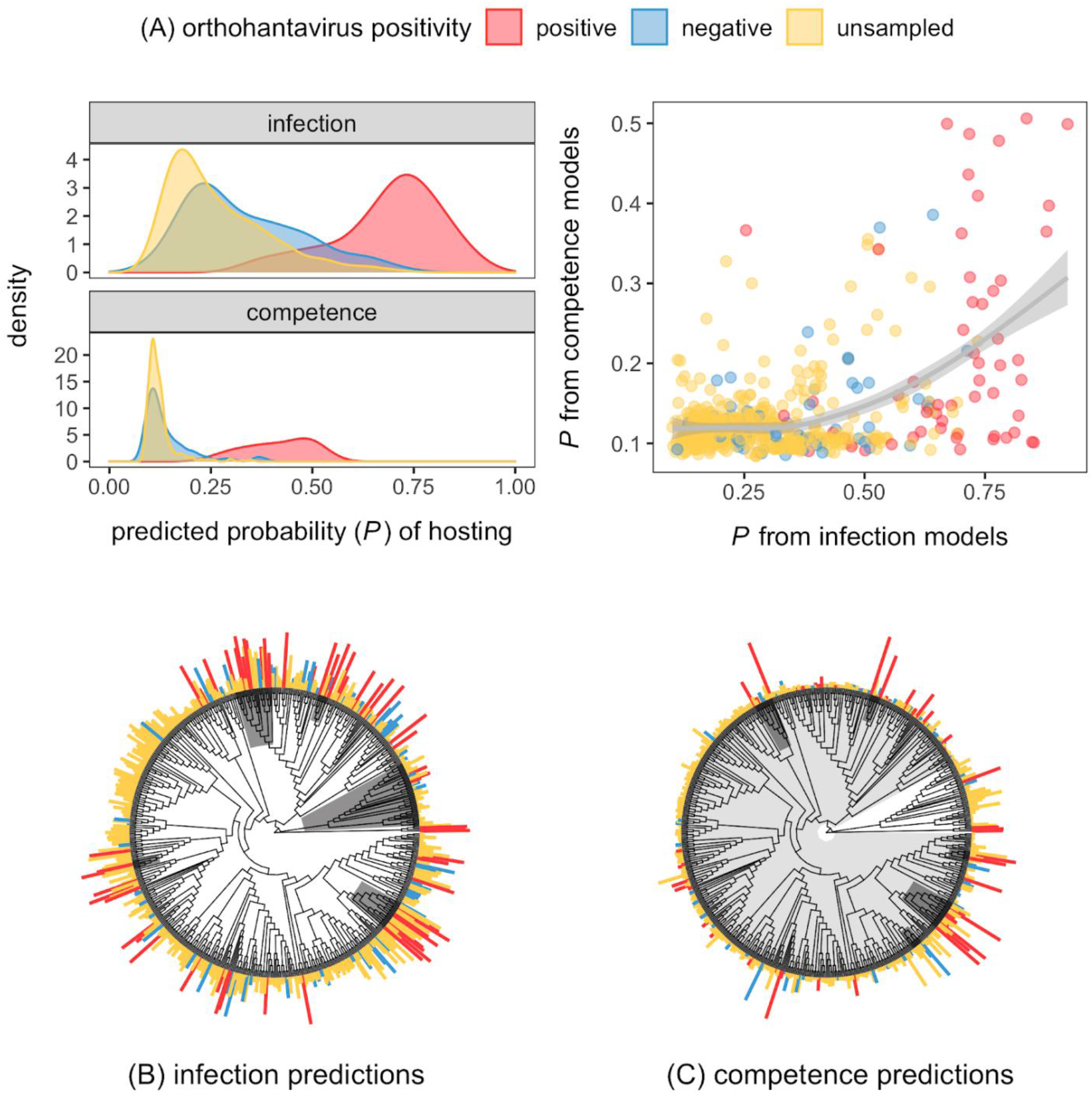
Predicted probabilities of rodent orthohantavirus positivity based on infection and competence.. (A) Distribution of propensity scores stratified by known positive, currently negative, and unsampled species. The scatterplot between predictions includes a smoothed curve and confidence intervals from a generalized additive model. (B–C) Taxonomic patterns in predictions as identified through phylogenetic factorization. Segments are scaled by probabilities and colored as in A. Clades identified with significantly different mean predictions are shown in grey, and additional information (e.g., included taxa, species richness) is included in S3 Table.

Lastly, we stratified our results into binary predictions using a 95% sensitivity threshold (S4 Table). This revealed a total 98 likely undiscovered hosts based on infection models versus only 14 undiscovered hosts based on competence models, of which 8 were also predicted by the former (Table 1). Mapping the geographic distribution of undetected hosts alongside known orthohantavirus-positive rodent species revealed that while predictions from infection models largely recapitulated the distributions of known RT-PCR-positive species, competence models suggested novel hotspots of overlapping reservoirs in the northeastern United States and northern South America, particularly along the Andes Mountains (Fig 4).

**Table 1.**
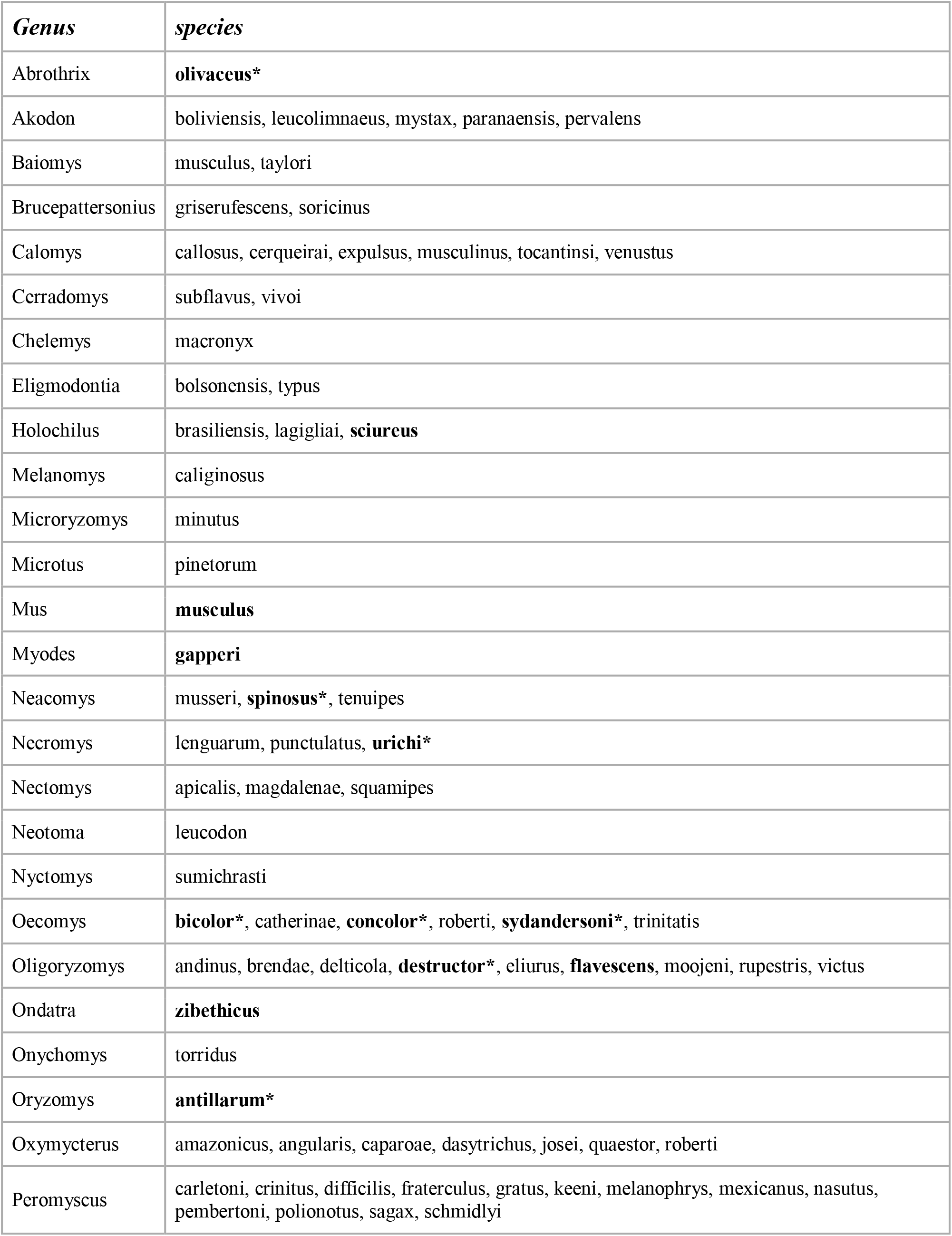
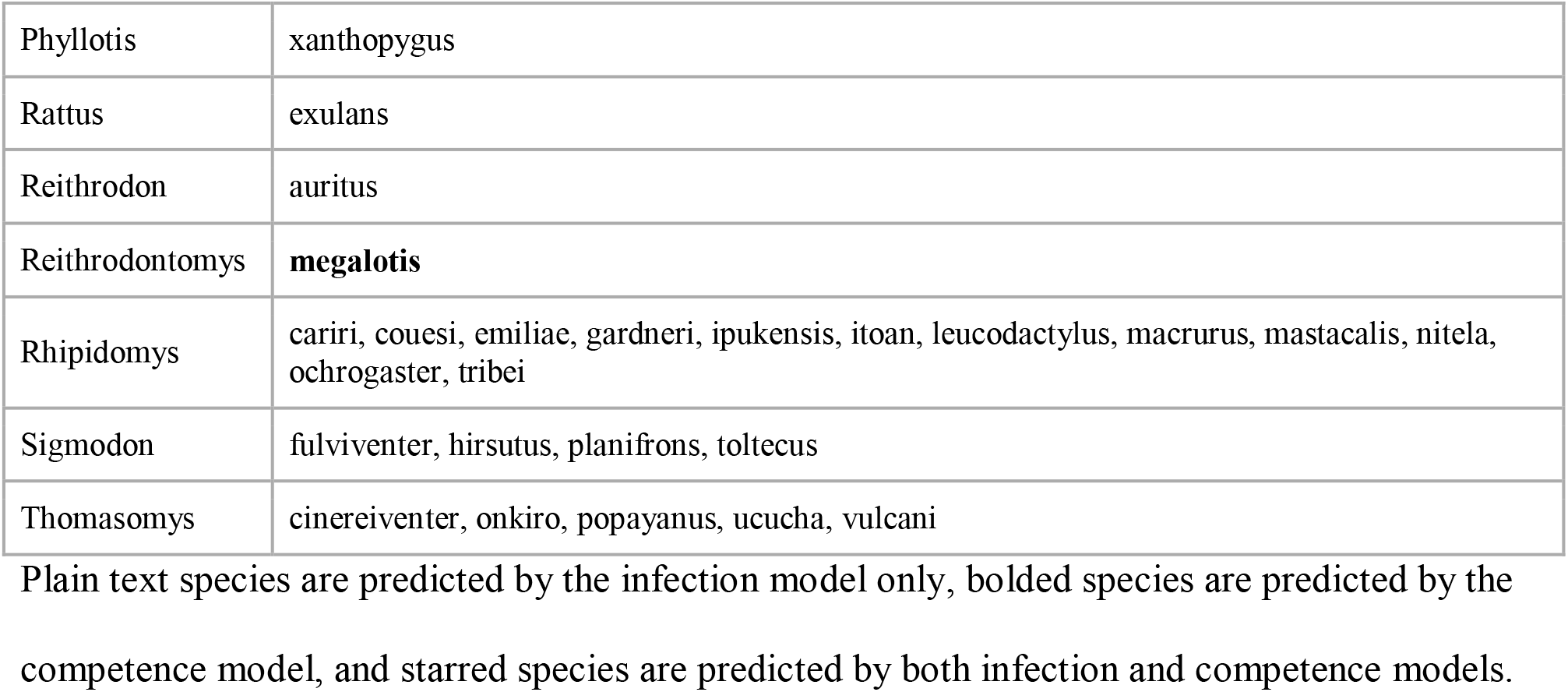
Predicted undiscovered hosts of hantaviruses: a priority list for future sampling efforts.

**Fig 4.**
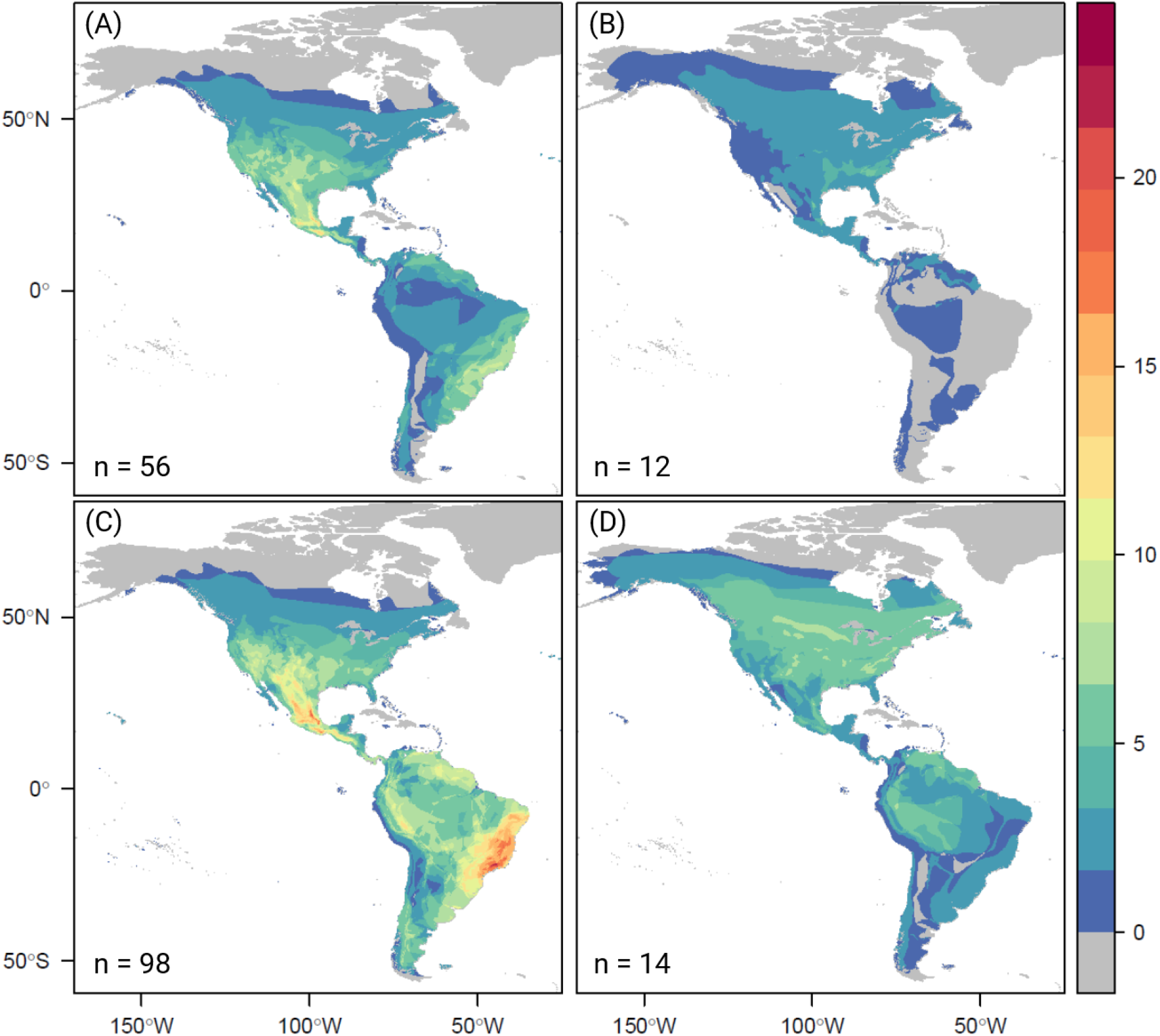
Distribution of orthohantavirus hosts. The distribution of known (A, B) and predicted undiscovered (C, D) hosts of orthohantaviruses based on infection (A,C) and competence (B, D).

Plain text species are predicted by the infection model only, bolded species are predicted by the competence model, and starred species are predicted by both infection and competence models.

We then mapped these total estimated sets of predicted unknown orthohantavirus hosts and reservoirs against anthropogenic impacts (Fig 5), as a proxy for cumulative and current excess spillover risk contributed by these environmental drivers. Hantavirus hosts coincide most with areas experiencing high anthropogenic impacts in central America and the Atlantic forests of Brazil and Uruguay. Reservoirs are distributed more evenly and extensively throughout the Americas, especially in both the Amazon and in high-latitude temperate ecosystems in North America. The greatest coincidence of those species with emergence risk factors may be in North American rural population centers and agricultural communities.

**Fig 5.**
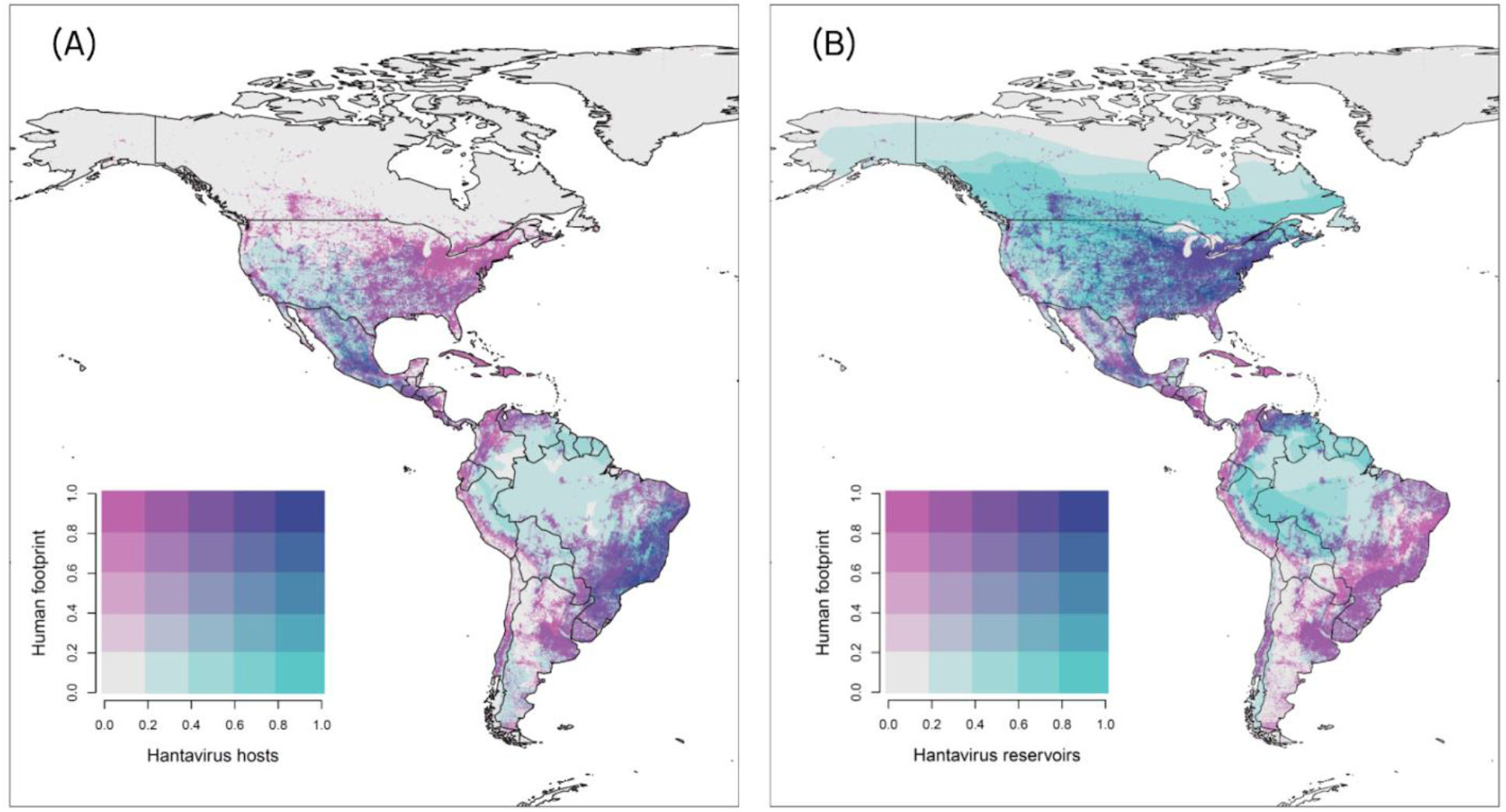
Geographic emergence risk of novel orthohantaviruses. Risk is presented as a function of total richness of predicted unknown rodent hosts (inferred from infection data; A) and reservoirs (inferred from competence; B) against the anthropogenic footprint on natural ecosystems.

## Discussion

In this study, we identified rodent species that are likely to host orthohantaviruses and demonstrated that the inclusion of competence data improves both model performance and generates distinct predictions from traditional models using RT-PCR data (i.e., infection). Determining the reservoir host of a particular orthohantavirus can be challenging, as orthohantaviruses are difficult to isolate (Strandin et al. 2020) and, like many infectious diseases, infection prevalence varies drastically across space and time (Walsh et al. 2007; Vadell et al. 2011; Holsomback et al. 2013). However, predictive modeling enables the detection of novel hosts in the absence of field data and in turn facilitates targeted field surveillance that can ultimately be used to mitigate hazards posed by zoonotic viruses (Becker et al. 2019).

Orthohantaviruses have traditionally been considered to follow evolutionary cophylogenies with their hosts, with few cross-species infections denoting distinct lineages (Hjelle et al. 1995; Herbreteau et al. 2006; Song et al. 2007). However, the discovery of additional orthohantaviruses has since expanded the diversity of hosts and demonstrated host switches in their evolutionary history (Blasdell et al. 2011; Guo et al. 2013). Indeed, orthohantaviruses have been isolated from species among all four subfamilies of muroid rodents in the Americas. Within those subfamilies, orthohantaviruses have been isolated from seven genera, and the subset of hosts predicted by both models would expand this range by four additional genera (Table 1). Additionally, the contrasting phylogenetic patterns of infection and competence, alongside the differing importance of taxonomic predictors in our two models, suggests that many orthohantavirus host-switches among disparate species have occurred, and frequent viral sharing among related species likely helps to account for the clusters of closely related cophylogenies (Fig 1).

In our study, postulated reservoir hosts are mostly concentrated in two regions, northeastern United States and northern South America, but southern Mexico and eastern Brazil are regions of likely spillover (Fig 4). Interestingly, all of these regions coincide with geographical gaps in known orthohantavirus distribution (Guzmán et al. 2017). In particular, not only would the discovery of an orthohantavirus hosted by *Myodes gapperi* bridge a geographic gap between Russia and North America, but it would also bridge a phylogenetic gap between Eurasian and American viruses. Several *Microtus* species are the only arvicoline rodents currently known to host orthohantaviruses in the New World, despite a variety of arvicoline hosts in the Old World (Blasdell et al. 2011).

Competence models (virus isolation data) were both more accurate and precise in estimating reservoir hosts compared to infection models (RT-PCR data). Although model performance (AUC) for our infection models was high, the higher AUC of the competence models (despite fewer rodent species with live virus isolation records) indicates the importance of including stronger evidence for reservoir capacity in model performance (Fig 2; Becker et al. 2020b). Additionally, there was substantial overlap in predicted host species, but the competence model produced a more concise list of virus reservoir candidates (Table 1). For example, in the present study, *Peromyscus* demonstrate the tendency for certain taxa to be infected more often without being reservoirs (Figs 1 and 3). Considerations of such differences in model performance are crucial when focusing surveillance efforts on likely reservoirs, and adherence to predictions based on infection status alone could lead to wasteful sampling of misidentified species.

Although this study focused on New World orthohantaviruses to enable higher resolution results in this system, our modeling approach is transferable to many other systems. Old World orthohantaviruses represent the most obvious extension, particularly for regions with minimal surveillance, such as Africa, the eastern Mediterranean, and Southeast Asia (Herbreteau et al. 2006; Guo et al. 2013). However, other virus groups that pose a threat to human welfare would also benefit from predictive modeling. For example, the reservoir hosts, and likely virus diversity, of orthopoxviruses (e.g., cowpox virus, monkeypox virus) are still mostly unknown, despite common evidence of orthopoxvirus infection among a diverse assemblage of wildlife, particularly rodents (McInnes et al. 2006; Kinnunen et al. 2011) and carnivores (Emerson et al. 2009; Morgan et al. 2019). In cases like this, models incorporating multiple levels of infection evidence can help filter out sampling “noise” to empower host detection for the many known and future emerging infectious diseases (Jones et al. 2008).

Predicted species in this study represent priority targets for orthohantavirus surveillance, particularly *Myodes gapperi* in North America and several members of the tribe Oryzomyini in South America. However, field verification of these predictions will be necessary to ultimately determine the true diversity of novel reservoirs. More broadly, we demonstrate here that the inclusion of competence data strengthens trait-based predictive modeling, and tailoring models based on outcomes of field studies will further improve accuracy. These methods will increase efficiency in host surveillance not only for orthohantaviruses, but also for a range of other pathogens important in human and wildlife health.

## Methods

### Hantavirus data

A systematic literature search was conducted in Web of Science to identify empirical studies that reported orthohantavirus infections in New World muroid rodents via RT-PCR (specific to negative-sense RNA viruses) or virus isolation (S1-3 Appendix). We recorded the number of studies per rodent species with each of the following criteria: at least one individual RT-PCR-positive; all individuals RT-PCR-negative; or virus isolation from at least one individual. Because orthohantaviruses cause persistent and chronic infections in rodents (Forbes et al. 2018), serological tests are often used to demonstrate current or recent infection, and RT-PCR is performed only on samples from antibody-positive individuals for virus characterization (Vaheri et al. 2008). To preclude false positives in these studies, only rodents that had positive RT-PCR results were considered RT-PCR-positive, and all other individuals were considered RT-PCR-negative, even if RT-PCR was not conducted. If a study used only serology without either RT-PCR or live virus isolation attempts, then the study was not included. When studies attempted virus isolation, additional RT-PCR results were recorded for specimen tissue analyses, but not infected cell cultures.

In studies that employed archived samples reported in a previous study (for the same level of evidence), those samples were omitted from our tallies to preclude pseudoreplication; instead, the original study was used. If a subsequent study examined a different level of evidence (e.g. virus isolation vs. RT-PCR), then we treated the two studies as a single report. In instances where the number or description of positive and negative results for each species was not clear in an article (including specimens reported at the genus level and outdated taxonomy that now represents multiple species), only definitive results were recorded. We manually matched select rodent species names between our orthohantavirus data and our phylogeny and trait data (see below). Species synonyms are provided in our online data repository. Since several *Rattus* and *Mus* are abundant in the Old and New World, only results derived in the Americas were included. Species without published evidence of orthohantavirus infection or competence were assigned pseudoabsences (Becker et al. 2020a).

### Phylogenetic analyses

We used a recently developed supertree of extant mammals to capture rodent phylogeny (Upham et al. 2019). The tree was simplified to our specified rodent species using the *ape* package in R (Paradis et al. 2004). Prior to predictive models, we conducted two assessments of phylogenetic signal (i.e., the propensity for related rodent species to be more similar in virus positivity). For both response variables (infection and competence), we used the *caper* package to calculate *D*, where a value of 1 indicates a phylogenetically random trait distribution and a value of 0 indicates phylogenetic clustering under a Brownian motion model of evolution (Fritz and Purvis 2010). Significant departure from either model was quantified using a randomization test with 1,000 permutations. However, because traits may also arise under a punctuated equilibrium model of evolution, we next used a graph-partitioning algorithm, phylogenetic factorization, to flexibly identify clades with significantly different propensity to be infected or competent at various taxonomic depths (Washburne et al. 2019). We used the *phylofactor* package to partition both outcomes as Bernoulli-distributed response variables with generalized linear models. We determined the number of significant phylogenetic factors (clades) using a Holm’s sequentially rejective 5% cutoff for the family-wise error rate.

### Rodent traits

We used a published dataset of 55 traits describing the morphology, geography, taxonomy, and life history of rodent species. Trait data were primarily from PanTHERIA alongside derived covariates including postnatal growth rate, relative age to sexual maturity, relative age at first birth, production, and species density (Jones et al. 2009; Han et al. 2015). We also used the *picante* package to quantify evolutionary distinctiveness, a measure of how isolated a species is within our muroid phylogeny (Kembel et al. 2010; Redding and Mooers 2006). Finally, we included binary covariates for our muroid rodent genera to represent taxonomy. We excluded predictors with no variance or missing values for over 75% of species, resulting in a total set of 62 biological covariates (S1 Table). Lastly, we used the *easyPubMed* package (accessed September 2020) to obtain the number of citations per species as a proxy for sampling effort (Olival et al. 2017; Fantini 2019).

### Boosted regression trees

We used boosted regression trees (BRTs) to classify rodent species as orthohantavirus hosts based on our predictor matrix of traits. BRTs circumvent many statistical issues associated with traditional hypothesis testing (e.g., a large number of predictors, complex interactions, non-randomly missing covariates) and can uncover new and surprising patterns in data to help develop testable hypotheses or predictions (Hochachka et al. 2007). Using this machine learning approach, we modeled binomial virus positivity separately for infection and competence.

BRTs maximize classification accuracy by learning patterns of features that best distinguish positive and negative hosts (Elith et al. 2008). This generates recursive binary splits for randomly sampled predictor variables, and successive trees are built using residuals of the prior best-performing tree as the new response. Boosting generates an ensemble of linked trees, where each achieves increasingly more accurate classification. Prior to analysis, we randomly split data into training (70%) and test (30%) sets while preserving the proportion of positive labels using the *rsample* package. Models were then trained with the *gbm* package (Greenwell et al. 2020), with the maximum number of trees set to 10000, a learning rate of 0.01, and an interaction depth of three. BRTs used a Bernoulli error distribution and five-fold cross-validation, and we used the *ROCR* package to quantify accuracy as area under the receiver operator curve (AUC; Sing et al. 2005). As results can depend on random splits between training and test data, we used 100 partitions to generate an ensemble (Evans et al. 2017). To diagnose if trait profiles of positive species are driven by study effort, we ran a secondary set of BRTs that modeled citation counts as a Poisson response (Plowright et al. 2019).

### Model performance and prediction

To assess how BRT performance varied between infection and competence models (Becker et al. 2020b), we used a paired *t*-test to compare AUC. We also assessed similarity in variable importance between models by estimating the Spearman correlation coefficient between feature ranks. Next, we predicted the probability of a species being positive for either response. When predicting species status, we set citation counts per species to their mean across species as a *post hoc* method to correct for sampling effort and remove at least some bias (Becker et al. 2020a). Lastly, we also estimated the Spearman correlation coefficients for the mean predictions between infection and competence models.

We used these mean predictions to identify “false negative” orthohantavirus hosts (i.e., those without a prior recorded orthohantavirus infection or isolation). We identified taxonomic patterns in predictions using Pagel’s λ as an estimate of phylogenetic signal with the *caper* package (Orme et al. 2013) as well as a secondary phylogenetic factorization to identify clades with significantly different predicted probabilities. To identify potential unknown hosts or reservoirs, we estimated a 95% sensitivity threshold using the *presenceabsence* package (Freeman and Moisen 2008), which can stratify predictions at a 5% omission rate on known true positives. This threshold, while fairly inclusive, mostly selects species with comparable probabilities of being infected or competent to known hosts.

To visualize the spatial distribution of known and predicted rodent hosts, we used the IUCN Red List database of mammal geographic ranges and overlaid these shapefiles for thresholded species based on infection and competence models. We finally mapped the distribution of known and predicted hosts and reservoirs against a proxy for cumulative anthropogenic impact on natural systems, given by the SEDAC Last of the Wild database’s 2009 Human Footprint map (Venter et al. 2016; Venter et al. 2018). This qualitative descriptor encompasses several geospatial layers that describe anthropogenic impacts with relevance to human exposure to rodents and hantaviruses, particularly human occupation (i.e., built-up settlements and human population), agricultural intensification (i.e., crop lands and pasture lands), and ecosystem fragmentation (i.e., road and railway density).

## Acknowledgements

This work was supported by the Viral Emergence Research Initiative (VERENA) consortium

## Supporting information

**S1 Appendix: Web of Science search terms for empirical study inclusion**. The focused search includes New World orthohantavirus names and abbreviations along with several terms for PCR and virus isolation. A separate non-focused search was also conducted that did not include the PCR and virus isolation terms.

**S2 Appendix: PRISMA diagram for empirical study inclusion**.

**S3 Appendix: Reference list for empirical studies used in analyses**.

**S1 Table. Feature coverage across the 601 muroid rodent species included in the BRT models**. Variables are presented as given in their original sources (Jones et al. 2009; Han et al. 2015).

**S2 Table. Rodent trait importance and ranks for BRTs trained on infection and competence**.

**S3 Table. Phylogenetic factorization of mean predicted probabilities for orthohantavirus positivity for (*i*) infection and (*ii*) competence models**. The number of retained clades after a 5% family-wise error rate, taxa corresponding to those clades, number of species per clade, and mean predicted probabilities for the clade compared to the paraphyletic remainder are shown.

**S4 Table. Sensitivity of estimated number of undiscovered hosts to thresholding method**.

**S1 Figure. Partial dependence plots for top traits for infection (left) and competence (right)**.

## Notes

### Competing Interest Statement

The authors have declared no competing interest.

