## Supplementary figures and images for "Viral competence data improves rodent reservoir predictions for American orthohantaviruses"

### Supplemental Appendix 2

**S2 Appendix: PRISMA diagram for empirical study inclusion.**

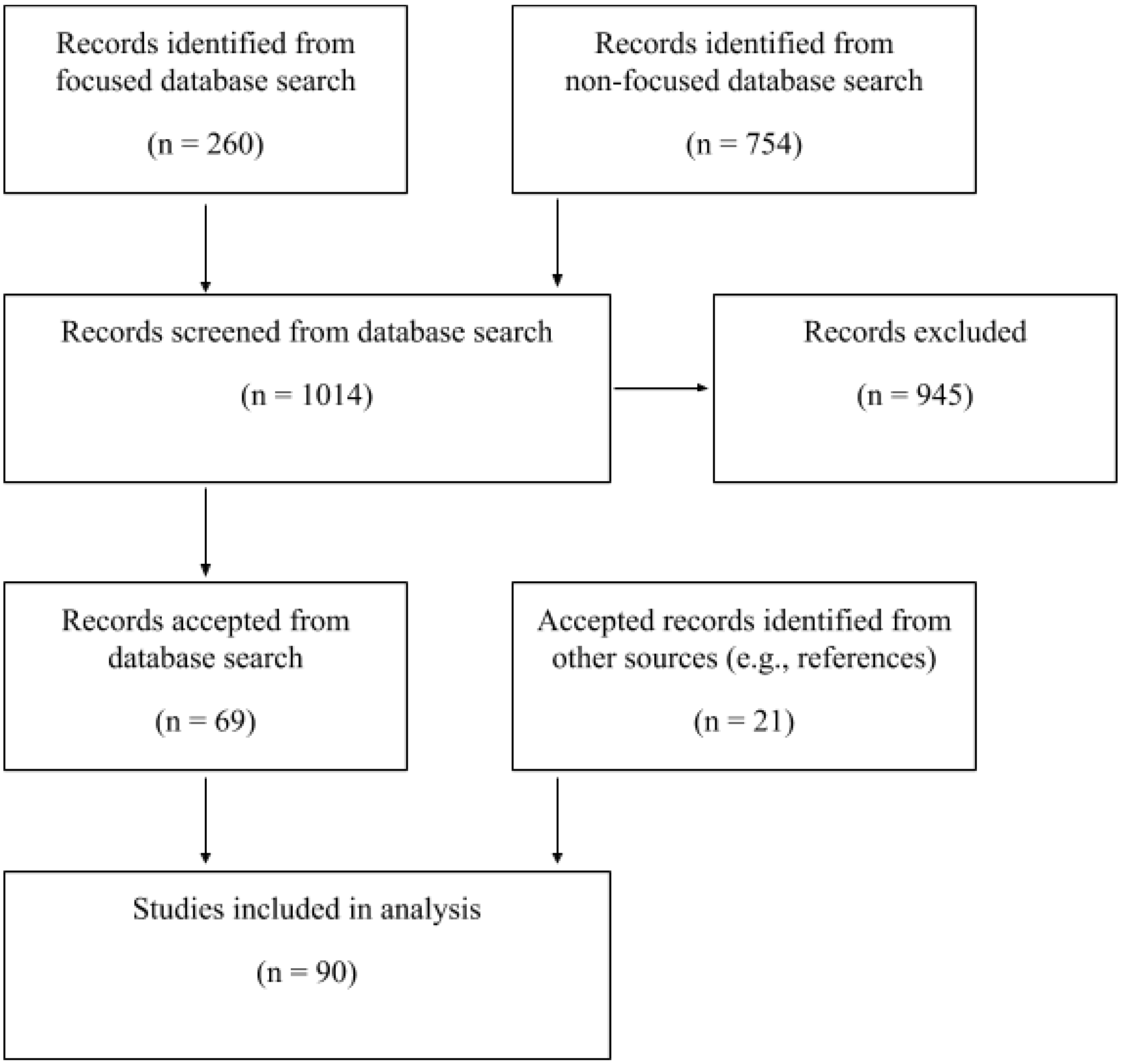

### Supplemental Figure 1

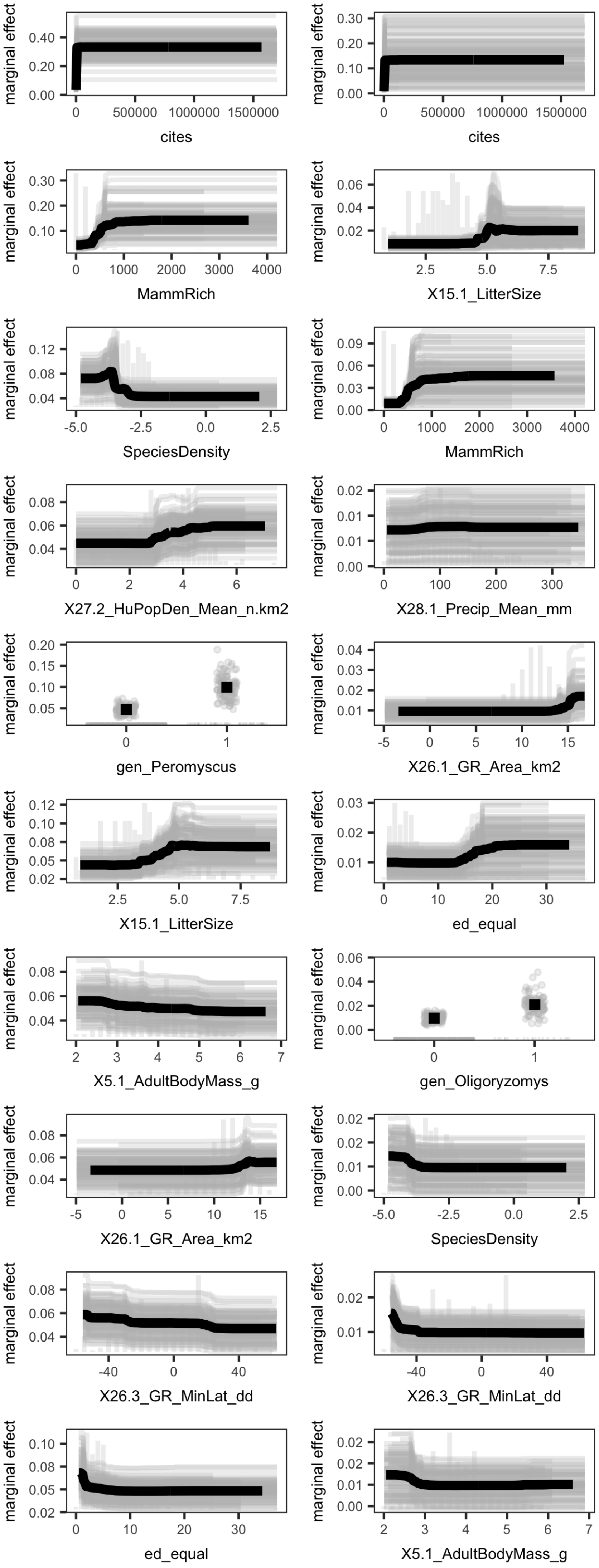
