## Supplemental Appendix 1 for "Viral competence data improves rodent reservoir predictions for American orthohantaviruses"

**S1 Appendix: Web of Science search terms for empirical study inclusion.** The focused search includes New World orthohantavirus names and abbreviations along with several terms for PCR and virus isolation. A separate non-focused search was also conducted that did not include the PCR and virus isolation terms.

Focused search string: TS=("Andes virus" OR "ANDV" OR "Araraquara virus" OR "ARAQV" OR "Juquitiba virus" OR "JUQV" OR "Maciel virus" OR "MACV" OR "Pergamino virus" OR "PRV" OR "PRGV" OR "Castelo dos Sonhos virus" OR "CASV" OR "Lechiguanas virus" OR "LECHV" OR "LECV" OR "Bermejo virus" OR "BMJV" OR "Oran virus" OR "ORNV" OR "Bayou virus" OR "BAYV" OR "Catacamas virus" OR "CATV" OR "Playa de Oro virus" OR "OROV" OR "Black Creek Canal virus" OR "BCCV" OR "Muleshoe virus" OR "MULV" OR "Cano Delgadito virus" OR "CADV" OR "Choclo virus" OR "CHOV" OR "Jabora virus" OR "JABV" OR "Carrizal virus" OR "CARV" OR "El Moro Canyon virus" OR "ELMCV" OR "Rio Segundo virus" OR "RIOSV" OR "Huitzilac virus" OR "HUIV" OR "Laguna Negra virus" OR "LANV" OR "Maripa virus" OR "MARV" OR "Rio Mamore virus" OR "RIOMV" OR "RMV" OR "Anajatuba virus" OR "ANAJV" OR "Rio Mearim virus" OR "RIOMMV" OR "Maporal virus" OR "MAPV" OR "Montano virus" OR "MNTV" OR "Necocli virus" OR "NECV" OR "Calabazo virus" OR "Prospect Hill virus" OR "PHV" OR "Isla Vista virus" OR "ISLAV" OR "Bloodland Lake virus" OR "BLV" OR "BLLLV" OR "ILV" OR "New York virus" OR "NYV" OR "Monongahela virus" OR "MGLV" OR "Blue River virus" OR "BRV" OR "Sin Nombre virus" OR "SNV" OR "FCV" OR "Limestone Canyon virus" OR "LCV" OR "Convict Creek virus" OR "CCV" OR "Muerto Canyon virus" OR "Seoul virus" OR "SEOV") AND TS=("PCR" OR "RT-PCR" OR "qPCR" OR "*PCR" OR "isolat*" OR "chain reaction" OR "extrac*") AND TS=("hantavirus" OR "orthohantavirus" OR "hantaviridae")
