## Supplemental Table 1 for "Viral competence data improves rodent reservoir predictions for American orthohantaviruses"

**S1 Table. Feature coverage across the 601 muroid rodent species included in the BRT models.**

| Feature | Coverage |
| --- | --- |
| cites | 1 |
| ed_equal | 1 |
| gen_Abrothrix | 1 |
| gen_Akodon | 1 |
| gen_Brucepattersonius | 1 |
| gen_Calomys | 1 |
| gen_Cerradomys | 1 |
| gen_Delomys | 1 |
| gen_Dicrostonyx | 1 |
| gen_Eligmodontia | 1 |
| gen_Euneomys | 1 |
| gen_Euryoryzomys | 1 |
| gen_Graomys | 1 |
| gen_Habromys | 1 |
| gen_Handleyomys | 1 |
| gen_Holochilus | 1 |
| gen_Hylaeamys | 1 |
| gen_Ichthyomys | 1 |
| gen_Microtus | 1 |
| gen_Neacomys | 1 |
| gen_Necromys | 1 |

| | |
|---|---|
| gen_Nectomys | 1 |
| gen_Neotoma | 1 |
| gen_Nephelomys | 1 |
| gen_Nesoryzomys | 1 |
| gen_Neusticomys | 1 |
| gen_Oecomys | 1 |
| gen_Oligoryzomys | 1 |
| gen_Oryzomys | 1 |
| gen_Oxymycterus | 1 |
| gen_Peromyscus | 1 |
| gen_Phyllotis | 1 |
| gen_Reithrodontomys | 1 |
| gen_Rheomys | 1 |
| gen_Rhipidomys | 1 |
| gen_Sigmodon | 1 |
| gen_Thomasomys | 1 |
| gen_Tylomys | 1 |
| MammRich | 0.9 |
| SpeciesDensity | 0.71 |
| X1.1_ActivityCycle | 0.32 |
| X12.1_HabitatBreadth | 0.33 |
| X12.2_Terrestriality | 0.32 |
| X15.1_LitterSize | 0.32 |

| | |
|---|---|
| X26.1_GR_Area_km2 | 0.77 |
| X26.2_GR_MaxLat_dd | 0.77 |
| X26.3_GR_MinLat_dd | 0.77 |
| X26.4_GR_MidRangeLat_dd | 0.77 |
| X26.5_GR_MaxLong_dd | 0.77 |
| X26.6_GR_MinLong_dd | 0.77 |
| X26.7_GR_MidRangeLong_dd | 0.77 |
| X27.1_HuPopDen_Min_n.km2 | 0.77 |
| X27.2_HuPopDen_Mean_n.km2 | 0.77 |
| X27.3_HuPopDen_5p_n.km2 | 0.77 |
| X27.4_HuPopDen_Change | 0.76 |
| X28.1_Precip_Mean_mm | 0.75 |
| X28.2_Temp_Mean_01degC | 0.75 |
| X30.1_AET_Mean_mm | 0.73 |
| X30.2_PET_Mean_mm | 0.73 |
| X5.1_AdultBodyMass_g | 0.75 |
| X6.1_DietBreadth | 0.26 |
| X6.2_TrophicLevel | 0.26 |

Variables are presented as given in their original sources (Jones et al. 2009; Han et al. 2015).
