## Supplemental Table 2 for "Viral competence data improves rodent reservoir predictions for American orthohantaviruses"

**S2 Table. Rodent trait importance and ranks for BRTs trained on infection and competence.**

| | Infection | | Competence | |
|---|---|---|---|---|
| **Feature** | **Importance** | **Rank** | **Importance** | **Rank** |
| cites | 0.273613 | 0.328172 | 1 | 1 |
| ed_equal | 0.032282 | 0.043723 | 10 | 6 |
| gen_Abrothrix | 1.00E-06 | 2.00E-06 | 41 | 35 |
| gen_Akodon | 2.90E-05 | 0 | 35 | 36 |
| gen_Brucepattersonius | 0 | 0 | 43 | 37 |
| gen_Calomys | 0.001407 | 0.003483 | 31 | 25 |
| gen_Cerradomys | 0 | 0 | 44 | 38 |
| gen_Delomys | 0 | 0 | 45 | 39 |
| gen_Dicrostonyx | 0 | 0 | 46 | 40 |
| gen_Eligmodontia | 0 | 0 | 47 | 41 |
| gen_Euneomys | 0 | 0 | 48 | 42 |
| gen_Euryoryzomys | 0 | 0 | 49 | 43 |
| gen_Graomys | 0 | 0 | 50 | 44 |
| gen_Habromys | 0 | 0 | 51 | 45 |
| gen_Handleyomys | 1.00E-06 | 0 | 42 | 46 |
| gen_Holochilus | 0 | 0 | 52 | 47 |
| gen_Hylaeamys | 0 | 0 | 53 | 48 |
| gen_Ichthyomys | 0 | 0 | 54 | 49 |
| gen_Microtus | 2.50E-05 | 6.90E-05 | 37 | 31 |
| gen_Neacomys | 0 | 0 | 55 | 50 |

| | | | | |
|---|---|---|---|---|
| gen_Necromys | 9.60E-05 | 0 | 33 | 51 |
| gen_Nectomys | 6.00E-06 | 0 | 39 | 52 |
| gen_Neotoma | 1.70E-05 | 0 | 38 | 53 |
| gen_Nephelomys | 0 | 0 | 56 | 54 |
| gen_Nesoryzomys | 0 | 0 | 57 | 55 |
| gen_Neusticomys | 0 | 0 | 58 | 56 |
| gen_Oecomys | 2.90E-05 | 8.00E-06 | 36 | 33 |
| gen_Oligoryzomys | 0.023508 | 0.03899 | 11 | 7 |
| gen_Oryzomys | 5.00E-05 | 0.018649 | 34 | 12 |
| gen_Oxymycterus | 0.005306 | 0 | 28 | 57 |
| gen_Peromyscus | 0.047574 | 4.60E-05 | 5 | 32 |
| gen_Phyllotis | 0 | 0 | 59 | 58 |
| gen_Reithrodontomys | 0.000389 | 4.00E-06 | 32 | 34 |
| gen_Rheomys | 0 | 0 | 60 | 59 |
| gen_Rhipidomys | 2.00E-06 | 0 | 40 | 60 |
| gen_Sigmodon | 0.003447 | 0.008429 | 30 | 20 |
| gen_Thomasomys | 0 | 0 | 61 | 61 |
| gen_Tylomys | 0 | 0 | 62 | 62 |
| MammRich | 0.110402 | 0.105545 | 2 | 3 |
| SpeciesDensity | 0.075615 | 0.033624 | 3 | 8 |
| X1.1_ActivityCycle | 0.017077 | 0.001947 | 15 | 30 |
| X12.1_HabitatBreadth | 0.005588 | 0.002467 | 27 | 27 |
| X12.2_Terrestriality | 0.011134 | 0.007967 | 22 | 21 |

| | | | | |
|---|---|---|---|---|
| X15.1_LitterSize | 0.047303 | 0.12484 | 6 | 2 |
| X26.1_GR_Area_km2 | 0.035525 | 0.049562 | 8 | 5 |
| X26.2_GR_MaxLat_dd | 0.009714 | 0.008787 | 25 | 19 |
| X26.3_GR_MinLat_dd | 0.035093 | 0.026583 | 9 | 9 |
| X26.4_GR_MidRangeLat_dd | 0.010228 | 0.002527 | 24 | 26 |
| X26.5_GR_MaxLong_dd | 0.01771 | 0.021711 | 14 | 11 |
| X26.6_GR_MinLong_dd | 0.018125 | 0.012047 | 13 | 16 |
| X26.7_GR_MidRangeLong_dd | 0.011001 | 0.006102 | 23 | 22 |
| X27.1_HuPopDen_Min_n.km2 | 0.0041 | 0.002248 | 29 | 28 |
| X27.2_HuPopDen_Mean_n.km2 | 0.056846 | 0.017936 | 4 | 13 |
| X27.3_HuPopDen_5p_n.km2 | 0.005739 | 0.002018 | 26 | 29 |
| X27.4_HuPopDen_Change | 0.013845 | 0.011122 | 18 | 17 |
| X28.1_Precip_Mean_mm | 0.016103 | 0.051138 | 17 | 4 |
| X28.2_Temp_Mean_01degC | 0.01175 | 0.013801 | 19 | 15 |
| X30.1_AET_Mean_mm | 0.01153 | 0.009133 | 20 | 18 |
| X30.2_PET_Mean_mm | 0.016688 | 0.005716 | 16 | 23 |
| X5.1_AdultBodyMass_g | 0.036151 | 0.022713 | 7 | 10 |
| X6.1_DietBreadth | 0.023455 | 0.014521 | 12 | 14 |
| X6.2_TrophicLevel | 0.011472 | 0.004377 | 21 | 24 |
