## Supplemental Appendix 3 for "Viral competence data improves rodent reservoir predictions for American orthohantaviruses"

**S3 Appendix: Reference list for empirical studies used in analyses.**

Araujo J, Pereira A, Nardi MS, Henriques DA, Lautenschalager DA, Dutra LM, et al. Detection of hantaviruses in Brazilian rodents by SYBR-Green-based real-time RT-PCR. Arch Virol. 2011;156:1269–74.

Bagamian K, Towner J, Mills J, Kuenzi A. Increased detection of Sin Nombre hantavirus RNA in antibody-positive deer mice from Montana, USA: Evidence of male bias in RNA viremia. Viruses. 2013;5:2320–8.

Belmar-Lucero S, Godoy P, Ferrés M, Vial P, Palma RE. Range expansion of *Oligoryzomys longicaudatus* (Rodentia, Sigmodontinae) in Patagonian Chile, and first record of hantavirus in the region. Rev Chil Hist Nat. 2009;82:265–75.

Bennett SG, Webb JP. Hantavirus (Bunyaviridae) infections in rodents from Orange and San Diego Counties, California. Am J Trop Med Hyg. 1999;60:75–84.

Bharadwaj M, Botten J, Torrez-Martinez N, Hjelle B. Rio Mamore virus: Genetic characterization of a newly recognized hantavirus of the pygmy rice rat, *Oligoryzomys microtis*, from Bolivia. Am J Trop Med Hyg. 1997;57:368–74.

Black WC, Doty JB, Hughes MT, Beaty BJ, Calisher CH. Temporal and geographic evidence for evolution of Sin Nombre virus using molecular analyses of viral RNA from Colorado, New Mexico and Montana. Virol J. 2009;6:102.

Boone JD, Mcgwire KC, Otteson EW, Debaca RS, Kuhn EA, Jeor SCS. Infection dynamics of Sin Nombre virus after a widespread decline in host populations. Am J Trop Med Hyg. 2002;67:310–8

Borucki MK, Boone JD, Rowe JE, Bohlman MC, Kuhn EA, DeBaca R, et al. Role of maternal antibody in natural infection of *Peromyscus maniculatus* with Sin Nombre virus. J Virol. 2000;74:2426–9.

Burek KA, Rossi CA, LeDuc JW, Yuill TM. Serologic and virological evidence of a Prospect Hill-like Hantavirus in Wisconsin and Minnesota. Am J Trop Med Hyg. 1994;51:286–94.

Cantoni G, Padula P, Calderon G, Mills J, Herrero E, Sandoval P, et al. Seasonal variation in prevalence of antibody to hantaviruses in rodents from southern Argentina. Trop Med Int Health. 2001;6:811–6.

Childs JE, Ksiazek TG, Spiropoulou CF, Krebs JW, Morzunov S, Maupin GO, et al. Serologic and genetic identification of *Peromyscus maniculatus* as the primary rodent reservoir for a new hantavirus in the southwestern United States. J Infect Dis. 1994;169:1271–80.

Childs JE, Korch GW, Glass GE, Leduc JW, Shah KV. Epizootiology of hantavirus infections in Baltimore: Isolation of a virus from Norway rats, and characteristics of infected rat populations. Am J Epidemiol. 1987;126:55–68.

Chu Y-K, Goodin D, Owen RD, Koch D, Jonsson CB. Sympatry of 2 hantavirus strains, Paraguay, 2003–2007. Emerg Infect Dis. 2009;15:1977–1980.

Chu Y-K, Owen RD, Sánchez-Hernández C, Romero-Almaraz Ma de L, Jonsson CB. Genetic characterization and phylogeny of a hantavirus from western Mexico. Virus Res. 2008;131:180–8.

Colombo VC, Brignone J, Sen C, Previtali MA, Martin ML, Levis S, et al. Orthohantavirus genotype Lechiguanas in *Oligoryzomys nigripes* (Rodentia: Cricetidae): New evidence of host-switching. Acta Trop. 2019;191:133–8.

de Figueiredo GG, Borges AA, Campos GM, Machado AM, Saggioro FP, Sabino Júnior G dos S, et al. Diagnosis of hantavirus infection in humans and rodents in Ribeirão Preto, State of São Paulo, Brazil. Rev Soc Bras Med Trop. 2010;43:348–54.

Dearing MD, Mangione AM, Karasov H, Morzunov S. Prevalence of hantavirus in four species of *Neotoma* from Arizona and Utah. J Mammal. 1998;79:1254–9.

Delfraro A, Clara M, Tomé L, Achaval F, Levis S, Calderón G, et al. Yellow pygmy rice rat (*Oligoryzomys flavescens*) and hantavirus pulmonary syndrome in Uruguay. Emerg Infect Dis. 2003;9:846–52.

Delfraro A, Tomé L, D'Elía G, Clara M, Achával F, Russi JC, et al. Juquitiba-like hantavirus from 2 nonrelated rodent species, Uruguay. Emerg Infect Dis. 2008;14:1447–51.

Della Valle MG, Edelstein A, Miguel S, Martinez V, Cortez J, Cacace ML, et al. Andes virus associated with hantavirus pulmonary syndrome in northern Argentina and determination of the precise site of infection. Am J Trop Med Hyg. 2002;66:713–20.

Dietrich N, Pruden S, Ksiazek TG, Morzunov SP, Camp JW. A small-scale survey of hantavirus in mammals from Indiana. J Wildl Dis. 1997;33:818–22.

Figueiredo LTM, Moreli ML, de Sousa RLM, Borges AA, de Figueiredo GG, Machado AM, et al. Hantavirus pulmonary syndrome, Central Plateau, southeastern, and southern Brazil. Emerg Infect Dis. 2009;15:561–7.

Firth C, Tokarz R, Simith DB, Nunes MRT, Bhat M, Rosa EST, et al. Diversity and distribution of hantaviruses in South America. J Virol. 2012;86:13756–66.

Fulhorst CF, Cajimat MNB, Utrera A, Milazzo ML, Duno GM. Maporal virus, a hantavirus associated with the fulvous pygmy rice rat (*Oligoryzomys fulvescens*) in western Venezuela. Virus Res. 2004;104:139–44.

Fulhorst CF, Monroe MC, Salas RA, Duno G, Utrera A, Ksiazek TG, et al. Isolation, characterization and geographic distribution of Caño Delgadito virus, a newly discovered South American hantavirus (family Bunyaviridae). Virus Res. 1997;51:159–71.

Guterres A, de Oliveira R, Fernandes J, Strecht L, Casado F, Gomes de Oliveira F, et al. Characterization of Juquitiba birus in *Oligoryzomys fornesi* from Brazilian Cerrado. Viruses. 2014;6:1473–82.

Henderson WW, Monroe MC, St. Jeor SC, Thayer WP, Rowe JE, Peters CJ, et al. Naturally occurring Sin Nombre virus genetic reassortants. Virology. 1995;214:602–60.

Himsworth CG, Bai Y, Kosoy MY, Wood H, DiBernardo A, Lindsay R, et al. An investigation of *Bartonella* spp., *Rickettsia typhi* , and Seoul hantavirus in rats ( *Rattus* spp.) from an inner-city neighborhood of Vancouver, Canada: Is pathogen presence a reflection of global and local rat population structure? Vector Borne Zoonotic Dis. 2015;15:21–6.

Hjelle B, Chavez-Giles F, Torrez-Martinez N, Yates T, Sarisky J, Webb J, et al. Genetic identification of a novel hantavirus of the harvest mouse *Reithrodontomys megalotis*. J Virol. 1994;68:6751–54.

Hjelle B, Lee SW, Song W, Torrez-Martinez N, Song JW, Yanagihara R, et al. Molecular linkage of hantavirus pulmonary syndrome to the white-footed mouse, *Peromyscus leucopus*: Genetic characterization of the M genome of New York virus. J Virol. 1995;69:8137–41.

Hjelle B, Anderson B, Torrez-Martinez N, Song W, Gannon WL, Yates TL. Prevalence and geographic genetic variation of hantaviruses of New World harvest mice (*Reithrodontomys*): Identification of a divergent genotype from a Costa Rican *Reithrodontomys mexicanus*. Virology. 1995;207:452–9.

Hjelle B, Torrez-Martinez N, Koster FrederickT. Hantavirus pulmonary syndrome-related virus from Bolivia. Lancet. 1996;347:57.

Holsomback TS, McIntyre NE, Nisbett RA, Strauss RE, Chu Y-K, Abuzeineh AA, et al. Bayou virus detected in non-Oryzomyine rodent hosts: An assessment of habitat composition, reservoir community structure, and marsh rice rat social dynamics. J Vector Ecol. 2009;34:9–21.

Jay M, Hjelle B, Davis R, Ascher M, Baylies HN, Reilly K, et al. Occupational exposure leading to hantavirus pulmonary syndrome in a utility company employee. Clin Infect Dis. 1996;22:841–4.

Johnson AM, Bowen MD, Ksiazek TG, Williams RJ, Bryan RT, Mills JN, et al. Laguna Negra virus associated with HPS in western Paraguay and Bolivia. Virology. 1997;238:115–27.

Kariwa H, Yoshida H, Sánchez-Hernández C, Romero-Almaraz M de L, Almazán-Catalán JA, Ramos C, et al. Genetic diversity of hantaviruses in Mexico: Identification of three novel hantaviruses from Neotominae rodents. Virus Res. 2012;163:486–94.

Kerins JL, Koske SE, Kazmierczak J, Austin C, Gowdy K, Dibernardo A. Outbreak of Seoul virus among rats and rat owners – United States and Canada, 2017. MMWR Morb Mortal Wkly Rep 2018;67:131–4.

Khan AS, Gaviria M, Rollin PE, Hlady WG, Ksiazek TG, Armstrong LR, et al. Hantavirus pulmonary syndrome in Florida: Association with the newly identified Black Creek Canal virus. Am J Med. 1996;100:46–8.

Kjemtrup AM, Messenger S, Meza AM, Feiszli T, Yoshimizu MH, Padgett K, et al. New exposure location for hantavirus pulmonary syndrome case, California, USA, 2018. Emerg Infect Dis. 2019;25:1962–4.

Klein SL, Shone SM, Glass GE, Zink MC, Hinson ER. Wounding: The primary mode of Seoul virus transmission among male Norway rats. Am J Trop Med Hyg. 2004;70:310–7.

Ksiazek TG, Nichol ST, Mills JN, Groves MG, Wozniak A, McAdams S, et al. Isolation, genetic diversity, and geographic distribution of Bayou virus (Bunyaviridae: Hantavirus). Am J Trop Med Hyg. 1997;57:445–8.

Kuenzi AJ, Douglass RJ, Bond CW, Calisher CH, Mills JN. Long-term dynamics of Sin Nombre viral RNA and antibody in deer mice in Montana. J Wildl Dis. 2005;41:473–81.

LeDuc JW, Smith GA, Johnson KM. Hantaan-like viruses from domestic rats captured in the United States. Am J Trop Med Hyg. 1984;33:992–8.

LeDuc JW, Smith GA, Pinheiro FP, Vasconcelos PFC, Rosa EST, Maiztegui JI. Isolation of a Hantaan-related virus from Brazilian rats and serologic evidence of its widespread distribution in South America. Am J Trop Med Hyg. 1985;34:810–5.

Lee P-W, Amyx HL, Yanagihara R, Gajdusek DC, Goldgaber D, Gibbs CJ. Partial characterization of Prospect Hill virus isolated from meadow voles in the United States. J Infect Dis. 1985;152:826–9.

Levis S, Morzunov SP, Rowe JE, Enria D, Pini N, Calderon G, et al. Genetic diversity and epidemiology of hantaviruses in Argentina. J Infect Dis. 1998;177:529–38.

Levis S, Rowe JE, Morzunov S, Enria DA, Jeor SS. New hantaviruses causing hantavirus pulmonary syndrome in central Argentina. Lancet. 1997;349:998–9.

Limongi JE, Oliveira RC, Guterres A, Costa Neto SF, Fernandes J, Vicente LHB, et al. Hantavirus pulmonary syndrome and rodent reservoirs in the savanna-like biome of Brazil's southeastern region. Epidemiol Infect. 2016;144:1107–16.

Londoño AF, Díaz FJ, Agudelo-Flórez P, Levis S, Rodas JD. Genetic evidence of hantavirus infections in wild rodents from northwestern Colombia. Vector Borne Zoonotic Dis. 2011;11:701–8.

Lopes LdB, Guterres A, Rozental T, Carvalho de Oliveira R, Mares-Guia M, Fernandes J, et al. *Rickettsia bellii*, *Rickettsia amblyommii*, and Laguna Negra hantavirus in an Indian reserve in the Brazilian Amazon. Parasit Vectors. 2014;7:191.

Martinez-Valdebenito C, Calvo M, Vial C, Mansilla R, Marco C, Palma RE, et al. Person-to-person household and nosocomial transmission of Andes hantavirus, southern Chile, 2011. Emerg Infect Dis. 2014;20:1637–44.

Matheus S, Guitet S, de Thoisy B, Donato D, Clément L, Lavergne A, et al. Maripa hantavirus in French Guiana: Phylogenetic position and predicted spatial distribution of rodent hosts. Am J Trop Med Hyg. 2014;90:988–92.

Medina RA, Torres-Perez F, Galeno H, Navarrete M, Vial PA, Palma RE, et al. Ecology, genetic diversity, and phylogeographic structure of Andes virus in humans and rodents in Chile. JVI. 2009;83:2446–59.

Meissner JD, Rowe JE, Borucki MK, St. Jeor SC. Complete nucleotide sequence of a Chilean hantavirus. Virus Res. 2002;89:131–43.

Milazzo ML, Cajimat MNB, Hanson JD, Bradley RD, Quintana M, Sherman C, et al. Catacamas virus, a hantaviral species naturally associated with *Oryzomys couesi* (Coues' oryzomys) in Honduras. Am J Trop Med Hyg. 2006;75(5): 1003–10.

Milazzo ML, Cajimat MNB, Romo HE, Estrada-Franco JG, Iñiguez-Dávalos LI, Bradley RD, et al. Geographic distribution of hantaviruses associated with neotomine and sigmodontine rodents, Mexico. Emerg Infect Dis. 2012;18:571–6.

Milazzo ML, Cajimat MNB, Richter MH, Bradley RD, Fulhorst CF. Muleshoe virus and other hantaviruses associated with neotomine or sigmodontine rodents in Texas. Vector Borne Zoonotic Dis. 2017;17:720–9.

Milazzo ML, Duno G, Utrera A, Richter MH, Duno F, de Manzione N, et al. Natural host relationships of hantaviruses native to western Venezuela. Vector Borne Zoonotic Dis. 2010;10:605–11.

Monroe MC, Morzunov SP, Johnson AM, Bowen MD, Artsob H, Yates T, et al. Genetic diversity and distribution of *Peromyscus*-borne hantaviruses in North America. Emerg Infect Dis. 1999;5:75–86.

Moro de Sousa RL, Moreli ML, Borges AA, Campos GM, Livonesi MC, Figueiredo LTM, et al. Natural host relationships and genetic diversity of rodent-associated hantaviruses in southeastern Brazil. Intervirology. 2008;51:299–310.

Morzunov SP, Rowe JE, Ksiazek TG, Peters CJ, St. Jeor SC, Nichol ST. Genetic analysis of the diversity and origin of hantaviruses in *Peromyscus leucopus* mice in North America. J Virol. 1998;72:57–64.

Nerurkar VR, Song J-W, Song K-J, Nagle JW, Hjelle B, Jenison S, et al. Genetic evidence for a hantavirus enzootic in deer mice (*Peromyscus maniculatus*) captured a decade before the recognition of hantavirus pulmonary syndrome. Virology. 1994;204:563–8.

Netski D, Thran BH, St. Jeor SC. Sin Nombre virus pathogenesis in *Peromyscus maniculatus*. J Virol. 1999;73:585–91.

Padula P, Figueroa R, Navarrete M, Pizarro E, Cadiz R, Bellomo C, et al. Transmission study of Andes hantavirus infection in wild sigmodontine rodents. JVI. 2004;78:11972–9.

Pesapane R, Enge B, Roy A, Kelley R, Mabry K, Trainor BC, et al. A tale of two valleys: Disparity in Sin Nombre virus antibody reactivity between neighboring Mojave Desert communities. Vector Borne Zoonotic Dis. 2019;19:290–4.

Powers AM, Mercer DR, Watts DM, Guzman H, Fulhorst CF, Popov VL, et al. Isolation and genetic characterization of a hantavirus (Bunyaviridae: Hantavirus) from a rodent, *Oligoryzomys microtis* (Muridae), collected in northeastern Peru. Am J Trop Med Hyg. 1999;61:92–8.

Raboni SM, Hoffmann FG, Oliveira RC, Teixeira BR, Bonvicino CR, Stella V, et al. Phylogenetic characterization of hantaviruses from wild rodents and hantavirus pulmonary syndrome cases in the state of Parana (southern Brazil). J Gen Virol. 2009;90:2166–71.

Ravkov EV, Rollin PE, Ksiazek TG, Peters CJ, Nichol ST. Genetic and serologic analysis of Black Creek Canal virus and its association with human Disease and *Sigmodon hispidus* infection. Virology. 1995;210:482–9.

Rawlings JA, Torrez-Martinez N, Neill SU, Moore GM, Hicks BN, Pichuantes S, et al. Cocirculation of multiple hantaviruses in Texas, with characterization of the small (S) genome of a previously undescribed virus of cotton rats (*Sigmododon hispidus*). Am J Trop Med Hyg. 1996;55:672–9.

Rhodes LV, Huang C, Sanchez AJ, Nichol ST, Zaki SR, Ksiazek TG, et al. Hantavirus pulmonary syndrome associated with Monongahela Virus, Pennsylvania. Emerg Infect Dis. 2000;6:616–21.

Rollin PE, Ksiazek TG, Elliott LH, Ravkov EV, Martin ML, Morzunov S, et al. Isolation of Black Creek Canal virus, a new hantavirus from *Sigmodon hispidus* in Florida. J Med Virol. 1995;46:35–9.

Rosa EST, Mills JN, Padula PJ, Elkhoury MR, Ksiazek TG, Mendes WS, et al. Newly recognized hantaviruses associated with hantavirus pulmonary syndrome in northern Brazil: Partial genetic characterization of viruses and serologic implication of likely reservoirs. Vector Borne Zoonotic Dis. 2005;5:11–9.

Rowe JE, Ksiazek TG, Riolo J, St. Jeor SC, Nichol ST, Rollin PE, et al. Occurrence of hantavirus within the rodent population of northeastern California and Nevada. Am J Trop Med Hyg. 1996;54:127–33.

Rowe JE, St. Jeor SC, Riolo J, Otteson EW, Monroe MC, Henderson WW, et al. Coexistence of several novel hantaviruses in rodents indigenous to North America. Virology. 1995;213:122–30.

Saasa N, Sánchez-Hernández C, de Lourdes Romero-Almaraz M, Guerrero-Ibarra E, Almazán-Catalán A, Yoshida H, et al. Ecology of hantaviruses in Mexico: Genetic identification of rodent host species and spillover infection. Virus Res. 2012;168:88–96.

Safronetz D, Cross RW, Fischer ER, Feldmann H, Voss TG, Waffa B, et al. Old World hantaviruses in rodents in New Orleans, Louisiana. Am J Trop Med Hyg. 2014;90:897–901.

Safronetz D, Drebot MA, Artsob H, Cote T, Makowski K, Lindsay LR. Sin Nombre virus shedding patterns in naturally infected deer mice ( *Peromyscus maniculatus* ) in relation to duration of infection. Vector Borne Zoonotic Dis. 2008;8:97–100.

Sanchez AJ, Abbott KD, Nichol ST. Genetic identification and characterization of Limestone Canyon virus, a unique *Peromyscus*-borne hantavirus. Virology. 2001;286:345–53.

Schmaljohn AA, Li D, Negly DL, Bressler DS, Turell MJ, Korch GW, et al. Isolation and initial characterization of a newfound hantavirus from California. Virology. 1995;206:963–972.

Sinclair JR, Carroll DS, Montgomery JM, Pavlin B, McCombs K, Mills JN, et al. Two cases of hantavirus pulmonary syndrome in Randolph County, West Virginia: A coincidence of time and place? Am J Trop Med Hyg. 2007;76:438–42.

Song J-W, Baek LJ, Nagle JW, Schlitter D, Yanagihara R. Genetic and phylogenetic analyses of hantaviral sequences amplified from archival tissues of deer mice (*Peromyscus maniculatus nubiterrae*) captured in the eastern United States. Arch Virol. 1996;141:959–967.

Song J-W, Baek L-J, Gajdusek DC, Yanagihara R, Gavrilovskaya I, Luft B, et al. Isolation of pathogenic hantavirus from white-footed mouse (*Peromyscus leucopus*). Lancet. 1994;344:1637.

Song W, Torrez-Martinez N, Irwin W, Harrison FJ, Davis R, Ascher M, et al. Isla Vista virus: A genetically novel hantavirus of the California vole *Microtus californicus*. J Gen Virol. 1995;76:3195–9.

Suzuki A, Bisordi I, Levis S, Garcia J, Pereira LE, Souza RP, et al. Identifying rodent hantavirus reservoirs, Brazil. Emerg Infect Dis. 2004;10:2127–34.

Toro J, Vega JD, Khan AS, Mills JN, Padula P, Terry W, et al. An outbreak of hantavirus pulmonary syndrome, Chile, 1997. Emerg Infect Dis. 1998;4:687–94.

Torres-Pérez F, Wilson L, Collinge SK, Harmon H, Ray C, Medina RA, et al. Sin Nombre virus infection in field workers, Colorado, USA. Emerg Infect Dis. 2010;16:308–10.

Travassos da Rosa ES, Medeiros DBA, Nunes MRT, Simith DB, Pereira A de S, Elkhoury MR, et al. Pygmy rice rat as potential host of Castelo dos Sonhos hantavirus. Emerg Infect Dis. 2011;17:1527–30.

Tsai TF, Bauer SP, Sasso DR, Whitfield SG, McCormick JB, Caraway TC, et al. Serological and virological evidence of a Hantaan virus-related enzootic in the United States. J Infect Dis. 1985;152:126–36.

Vadell MV, Bellomo C, San Martín A, Padula P, Gómez Villafañe I. Hantavirus ecology in rodent populations in three protected areas of Argentina: Hantavirus ecology in protected areas. Trop Med Int Health. 2011;16:1342–52.

Vincent MJ, Quiroz E, Gracia F, Sanchez AJ, Ksiazek TG, Kitsutani PT, et al. Hantavirus pulmonary syndrome in Panama: Identification of novel hantaviruses and their likely reservoirs. Virology. 2000;277:14–9.

White DJ. Human and rodent hantavirus infection in New York state: Public health significance of an emerging infectious disease. Arch Intern Med. 1996;156:722–6.

Yee J, Wortman IA, Nofchissey RA, Goade D, Bennett SG, Webb JP, et al. Rapid and simple method for screening wild rodents for antibodies to Sin Nombre hantavirus. J Wildl Dis. 2003;39:271–7.
