## Supplemental Appendix 3 for "Viral competence data improves rodent reservoir predictions for American orthohantaviruses"

**S3 Table. Phylogenetic factorization of mean predicted probabilities for orthohantavirus positivity for (*i*) infection and (*ii*) competence models.**

| | Factor | Taxa | Species | Clade | Other |
|---|---|---|---|---|---|
| | 1 | Oligoryzomys | 20 | 0.61 | 0.3 |
| | 2 | Peromyscus_pembertoni, Peromyscus_interparietalis, Peromyscus_mekisturus, Peromyscus_crinitus, Peromyscus_dickeyi, Peromyscus_spicilegus, Peromyscus_winkelmanni, Peromyscus_aztecus, Peromyscus_hylocetes, Peromyscus_boylii, Peromyscus_simulus, Peromyscus_stephani, Peromyscus_madrensis, Peromyscus_beatae, Peromyscus_levipes, Peromyscus_carletoni, Peromyscus_schmidlyi, Peromyscus_eva, Peromyscus_furvus, Peromyscus_attwateri, Peromyscus_nasutus, Peromyscus_difficilis, Peromyscus_truei, Peromyscus_gratus, Peromyscus_ochraventer | 25 | 0.51 | 0.31 |
| | 3 | Oxymycterus_hispidus, Oxymycterus_quaestor, Oxymycterus_angularis, Oxymycterus_rufus, Oxymycterus_josei, Oxymycterus_caparoae | 6 | 0.58 | 0.31 |
| | 4 | Peromyscus_guardia, Peromyscus_polionotus, Peromyscus_maniculatus, Peromyscus_keeni, Peromyscus_melanotis, Peromyscus_leucopus, Peromyscus_gossypinus | 7 | 0.54 | 0.31 |
| (*i*) | 5 | Dicrostonyx, Phenacomys, Arborimus, Ondatra, Neofiber, Lemmus, Synaptomys, Lemmiscus, Microtus, Myodes | 43 | 0.2 | 0.32 |
| | 1 | Oligoryzomys | 20 | 0.23 | 0.13 |
| | 2 | Oryzomys_couesi, Oryzomys_nelsoni, Oryzomys_antillarum, Oryzomys_dimidiatus, Oryzomys_gorgasi, Oryzomys_palustris | 6 | 0.26 | 0.13 |
| | 3 | Peromyscus_guardia, Peromyscus_polionotus, Peromyscus_maniculatus, Peromyscus_keeni, Peromyscus_melanotis, Peromyscus_leucopus, Peromyscus_gossypinus | 7 | 0.22 | 0.13 |
| | 4 | Oecomys_bicolor, Oecomys_roberti, Oecomys_superans, Oecomys_trinitatis, Oecomys_mamorae, Oecomys_flavicans, Oecomys_sydandersoni | 7 | 0.22 | 0.13 |
| (*ii*) | 5 | Neotoma, Xenomys, Nelsonia, Hodomys, Ochrotomys, Reithrodontomys, Isthmomys, Onychomys, Osgoodomys, Peromyscus, Megadontomys, Habromys, Podomys, Neotomodon, Scotinomys, Baiomys, Tylomys, Ototylomys, Otonyctomys, Nyctomys, Sigmodon, Rheomys, Ichthyomys, Wilfredomys, | 512 | 0.12 | 0.19 |

| | | | | | |
|---|---|---|---|---|---|
| | | Anotomys, Reithrodon, Neusticomys, Chibchanomys, Phaenomys, Chinchillula, Abrawayaomys, Kunsia, Scapteromys, Akodon, Bibimys, Lenoxus, Blarinomys, Brucepattersonius, Thalpomys, Necromys, Podoxymys, Deltamys, Thaptomys, Oxymycterus, Juscelinomys, Rhipidomys, Thomasomys, Rhagomys, Chilomys, Aepeomys, Scolomys, Zygodontomys, Megaoryzomys, Hylaeamys, Mindomys, Nephelomys, Oecomys, Euryoryzomys, Transandinomys, Handleyomys, Neacomys, Microryzomys, Oreoryzomys, Microakodontomys, Holochilus, Pseudoryzomys, Amphinectomys, Nectomys, Nesoryzomys, Aegialomys, Sigmodontomys, Melanomys, Oryzomys, Noronhomys, Pennatomys, Megalomys, Lundomys, Cerradomys, Sooretamys, Drymoreomys, Eremoryzomys, Neotomys, Euneomys, Irenomys, Andinomys, Punomys, Delomys, Calomys, Phyllotis, Paralomys, Galenomys, Auliscomys, Loxodontomys, Tapecomys, Salinomys, Andalgalomys, Graomys, Eligmodontia, Calassomys, Wiedomys, Juliomys, Chelemys, Pearsonomys, Geoxus, Notiomys, Abrothrix | | | |
| | 6 | Sigmodon | 13 | 0.17 | 0.13 |

The number of retained clades after a 5% family-wise error rate, taxa corresponding to those clades, number of species per clade, and mean predicted probabilities for the clade compared to the paraphyletic remainder are shown.
