## Supplemental Table 4 for "Viral competence data improves rodent reservoir predictions for American orthohantaviruses"

**S4 Table. Sensitivity of estimated number of undiscovered hosts to thresholding method.**

| | PCR model | Isolation model |
|---|---|---|
| Maximum kappa statistic | 5 | 0 |
| Maximum percent correctly classified | 5 | 0 |
| 90% sensitivity (10% omission) | 48 | 8 |
| Equal sensitivity and specificity | 49 | 14 |
| 95% sensitivity (5% omission) | 98 | 14 |
